# Live-Exudation Assisted Phytobiome Cultromics System (LEAP-CS): A High-Throughput Cultromics System for Studying Plant-Microbiome Interactions through Diffusible Metabolic Exchange

**DOI:** 10.1101/2025.05.07.652788

**Authors:** Mrinmoy Mazumder, Shruti Pavagadhi, Raktim Bhattacharya, Arijit Mukherjee, Seyed Mohammad Majedi, Ivan Chin Hin Tan, Sanjay Swarup

## Abstract

This study introduces an innovative methodology for co-cultivating plants and microbes, employing a membrane-based structure to facilitate their physical separation. The Live-Exudation Assisted Phytobiome Cultromics System (LEAP-CS) has the capability to investigate the complex interaction of plant and soil microbiome under in vitro conditions, offering potential benefits. Subsequent validation and testing can be performed through pot, greenhouse, and field trials. The system can efficiently function as a high-throughput screening tool for assessing plant-microbiome interactions and their associated metabolic signatures. Our phytobiome culturing technique harnesses root exudation from live plants, capitalizing on the membrane’s selective separation to prevent direct physical interaction between plant and microbiome components. Consequently, the interaction is solely through chemical-mediated signalling. By employing this method, we can intricately dissect complex plant-microbiome interactions while faithfully maintaining and emulating the contact independent inherent associations prevalent within these phytobiomes. In conclusion, this user-friendly and reproducible ’in-vitro’ model holds immense potential for shedding light on the intricate community and metabolic exchange dynamics of plant-microbiome interactions, thus significantly advancing our understanding in this area.

## 1. Introduction

### 1.1 Background and motivation

Healthy plants are colonized by a rich diversity of microbes, forming complex microbial consortia that impact plant growth and productivity ^1,2^. These plant-associated microbiomes possess the requisite genetic machinery to regulate soil carbon stocks, cycle essential plant-nutrients, confer resistance to plants from invasive pathogenic microorganisms, and degrade pollutants ^3^. There is a scientific cognizance that utilizing the multidimensional benefits of these microbiomes can prove beneficial for sustainable agriculture by minimizing chemical and fertilizer inputs translating to lower carbon footprints ^4,5^. However, their wide use as adjuncts in current agricultural practices is largely restricted by our limited understanding of these complex plant-microbiome associations ^6,7^. Among the various ecological niches of plant-microbiomes, the rhizosphere, a narrow zone surrounding and influenced by plant roots, is considered as the hotbed for below-ground plant-microbiome associations ^8^. Notably, rhizosphere is regarded as one the most complex ecosystems on the earth ^9^. To this end, characterizing model rhizosphere microbial communities, which occupies the niche developed by the gradients of root exudates in the rhizosphere region ^10^ is of special interest due to its probable direct role in providing specific factors for plant growth. Consequently, these efforts can pave the way for developing and engineering beneficial plant-associated microbiomes for climate-resilient, nutrient-efficient, and sustainable food crops ^11, 12^. Recent trends in microbiomics have established robust methods to identify the potential or realized beneficial function by specific members of phytobiomes (microbial community associated with plants) along with the cross-kingdom chemical signalling between plants and microbes ^6,13^.Major gaps, however, remain in, (i) determining a systematic way to identify minimal set of the microbial community members that can provide desired services to the host crop plants; (ii) a high throughput *in vitro* system to test the effects of microbes or microbial solutions on plants before deploying them in green house or field conditions; (iii) ensuring system stability and active functionality of the microbes in context of metabolite exchange at the right stage of host crop; (iv) identifying low-cost and easy system to conduct research on plant-microbe interaction ^11,14,15^. Our rationale for addressing some of these gaps and challenges is to take the first step in bringing a large community into a cultured based format that relies on the host services for its assembly and growth. In the past, attempts have been made to culture microbial strains using specific substrates, metabolites as found in root exudates or soil extracts in media. As root exudation patterns are highly dynamic and depends on host species and growth environmental conditions, such root exudates only represent a limited scenario within a bigger range of exudates availability for microbiome associations ^16,17^.Another challenge in the field of host microbiome interactions has been to deconvolute the confounded effects of microbial attachment to surfaces and metabolite exchanges. Acknowledging the fact that millions of microbes provide host beneficial service in the rhizosphere which are located few millimetre distance away from the roots, highlights that we need to develop approaches to identify microbes that provide the service within narrow distance ranges.

To this end, having a system which can mimic many, but not all complexities of soil-based systems under *in vitro* conditions could be beneficial. Outcomes from such a system can then be further validated and tested in pot, green house and eventual field trials. Such a system can serve as an efficient high-throughput screening tool to measure plant-microbiome interactions and their typical metabolic signatures. Here, we describe a plant–microbiome mesocosm culturing system that physically separates the plant and microbial components, allowing only small molecules, such as root or microbial exudates, to pass between them. This system allows us to tease apart complex plant-microbiome interactions, while preserving and mimicking the natural associations within these phytobiomes in an easy and reproducible *“in-vitro”* format.

### 1.2 Development of the system

Against the light of the experimental challenges in the phytobiome domain to study plant-microbiome interactions, we developed a system driven by ecological principles that govern these interactions within the complex phytobiomes. Our system favours and allows for enhancement of natural ecosystem processes that support intricate, yet dynamic plant-microbiome associations. Our system is designed to (i) simulate native associations between the plant hosts and its associated microbiomes; (ii) allow metabolic exchange and chemical dialogue to enhance processes underlying plant-microbe interactions; (iii) enable concurrent data collection of plant phenotypic/molecular traits, microbiome composition, and interacting metabolites. To address the scientific needs of a wider scientific community, we made the system modular and scalable to an extent. Modularity of our system supports (i) easy manipulation of the microbiome communities; (ii) chemical complementation assays; (iii) different biological and analytical platforms for phenotyping; (iv) non-invasive *in-situ* growth-profiling. With the current features of our system, we can systematically capture (i) plant-microbe and microbe-microbe interactions; (ii) metabolic exchanges and crosstalk; (iii) microbial community shifts through simultaneous multi-omics analyses at different levels for integrated biological outputs. Consequently, our system can be utilized for both mechanistic and translational studies involving interactions within phytobiomes at various levels.

LEAP-CS has six modules that are assembled sequentially: 1. Plate-base lid; 2. Solid growth support media; 3. Layer of microbiome; 4. Membrane filter (preventing direct physical contact between host plants and its microbiome); 5. Plant seedlings; 6. Plate-base top. A schematic describing all the modules of the system along with the assembled system is provided in Figure 1.

**Figure 1:**
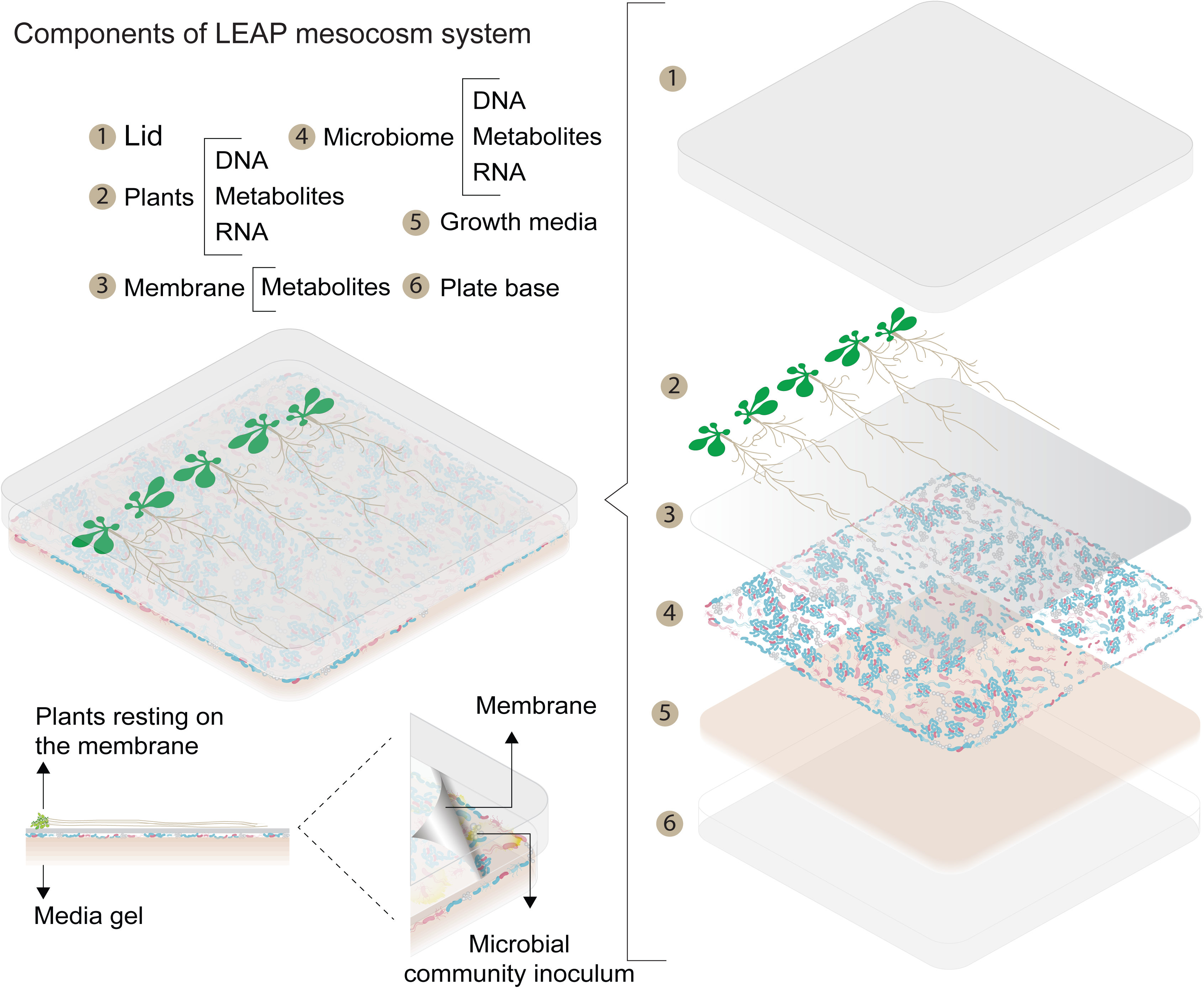
Overview of the Live Exudation-Assisted Phytobiome – Culturomics System (LEAP-CS) with its modular components. Module 1: plate lid; Module 2: Plant Hosts; Module 3: Membrane separating the plant hosts (#2) and Microbiome (#4); Module 4: Microbiome layer; Module 5: Growth Substrate/Media; Module 6: Plate base.

#### Module 1 and 6-Plate base/lid

We have utilized commonly available disposable plastic polystyrene petridish. We opted for rectangular dish (12 cm X12 cm X 1.7 cm, vented, Sigma-Aldrich, Missouri, USA) which is suitable for our host plants (*Arabidopsis Thaliana*). It is pertinent to note that these petri-dishes are available in variety of sizes, shapes and can be customized to achieve preferred dimensions to suit other host plants which may require longer assay run time and larger space to accommodate the seedlings. Owing to its widespread use, low-cost and durability we chose simple petri-dish to host this mesocosm assay set-up. There is some evidence that polystyrene material may result in some sample losses due to adherence of certain materials (for instance metabolites) as a result of its surface chemistry and physical properties. If this is of concern in the experimental design, then petridish made of other substrates like glass can also be considered which are compatible with our system. If experimental design involves addition of any chemical which is photolabile (light sensitive), then amber coloured petridishes may be considered, however, they may affect plant growth as they will block the incoming light.

#### Module 2-Solid support media

This module supports the growth of the host plants in the assay. For our set-up, we have tested some commonly used plant growth support media like murashige and skoog (MS) medium (without sucrose) water agar and soil extract agar. All of them could support the growth of host plants throughout the assay run time. Other kinds of plant growth media like linsmaier and skoog (LS) medium, gamborg (B5) medium and nitsch and nitsch (NN) medium can also be used with our system. A variety of gelling agents such as agar, agarose, gellan gum can be considered for the support media if it is compatible with the target host plants. Before adopting any growth medium for the system, its compatibility with host plants and microbiome should be established. Nutrient-rich growth media can sometimes overwhelm the system with microbial growth which also increase chances of cross-contamination. Customized growth media could be another alternative to conventional support media which can simulate specific environmental conditions for host plants and its associated microbiomes.

#### Module 3-layer of microbiome

This module is laid on the solid support media in the assay set-up. For establishing the protocol for LEAP-CS we used native microbiome from the host plants which was enriched *in-situ*, retrieved and then spread on the solid support media. Before applying the microbiome to the assay, we performed inoculum characterization through flow cytometry to evaluate the cell counts. Similar methods to characterize the inoculum should be employed before introducing it into the system as microbiome will influence the plant phenotypes as well as the chemical exchange through live exudation. Dose-dependent response should also be evaluated to determine the right amount of microbiome inoculum required in the assay. This step is critical to ensure reproducibility of the results obtained from this assay. Although, system is designed to study plant-microbiome interactions, layer of the microbiome can be replaced by a single microbial strain or any predetermined microbial community/synthetic consortia. However, inoculum characterization will still be required, and it should be optimized before the onset of the actual experiment. More details on this application are available elsewhere in the manuscript.

#### Module 4-membrane

This module sits on the lawn of microbiome in LEAP-CS and is composed of layer of thin cellulose dialysis membrane (commercially available, Sigma-Aldrich, Missouri, USA). Membrane excludes substances larger than 14 kDa to pass through, thus preventing direct contact between plants and microbiome, while still allowing for free and dynamic metabolic exchange between them. This membrane can be UV-sterilized prior to LEAP-CS assembly. In our system, minimal structural damage was observed in membrane after 10 days and we could retrieve the membrane with operational ease. However, if assay runs longer, some amount of membrane disintegration is possible which could be related to the interactions between the microbiome and membrane material. If the experiment calls for a longer assay run time, then membranes made of other materials with similar pore size can be considered which have higher resilience.

#### Module 5-Host Plants

Model plant species were utilized to optimize assay conditions. However, the system has also been tried on other plant hosts (Brassica, green leafy vegetables) and system conditions were compatible for these plant hosts. We believe assay run time and solid support media would need to be adapted for the host plant of interest. Selection of the host plant will largely depend on the experimental question; however, we strongly recommend optimizing the growth conditions of host plants in the assay prior to the actual experiment.

### 1.3 Comparison with other methods

As discussed in the earlier section, there are several systems available to study plant-microbe; plant-microbiome interactions. While those built on plant-microbiome models, are largely limited to hydroponic or soilless set-ups. We provide here a comparison for some of the representative methods (Table 1) available to the scientific community which are closest to the system described in this paper.

**Table 1:**

| Reference | System type | Characteristics |
| --- | --- | --- |
| Gao et al.,<br>2018<sup>18</sup> | Plant-microbe/plant-microbiome | + non-destructive imaging of root morphology and metabolite characterization using spatial imaging; provision to install LED to evaluate effect of light intensity on root growth |
|  |  | - Horizontal placement; gravitropic response cannot be simulated; system is not compatible with automated sampling; no provision to perform any other omics experiments as such; plant phenotyping is also difficult to achieve as system is restricted by number of plants; only crude characterization of microbiome is possible through the system |
|  |  | +Plant and microbe interactions can be captured in the system; system is |
| Nathoo et al., 2017 <sup>19</sup> | Plant-microbe | compatible for multi-omics like metabolomics, transcriptomics, genomics |
|  |  | -Hydroponic system so observed outcomes will be limited to interactions that are relevant for liquid nutrient media; system captures plant-microbe interactions which may not really reflect the complex interrelationships in phytobiome |
| Korenblam et al., 2020 <sup>20</sup> | Split root- Plant-microbe/plant-microbiome | +Captures systemic and localized effects caused by phytobiome interactions; system allows for evaluating both single microbe and complex microbiome communities; system allows to capture both root exudates as well as tissue metabolome |
|  |  | -system is not suitable for some plants with single tap root; hydroponic system so observed outcomes will be limited to interactions that are relevant for liquid nutrient media |
| Farag et al., 2017 <sup>21</sup> | Plant-microbe/microbe-microbe | +Evaluates effects of bacterial volatile compounds on plant growth, captures "no-contact" plant-microbe interactions; allows studying effect of volatiles on bacterial motility as well as antibiotic resistance |
|  |  | -Does not recapitulate interactions in natural conditions; system is suitable only to access plant-microbe and microbe-microbe interactions. |
|  |  | +Allows non-destructive repeated sampling as well as non-invasive imaging of root growth; Responsiveness to localized |
| Wei et al., 2019 <sup>22</sup> | Rhizobox- Plant-microbiome | nutrient patches can be monitored |
|  |  | -Construction of the system is time-consuming and maintaining homogeneous soil moisture is difficult. |
| Kremer et al., 2021 <sup>23</sup> | Peat-based gnotobiotic system- Plant-microbe/plant-microbiome | +Supports microbial growth due to peat-based substrates; Easily accessible, high-throughput and versatile |
|  |  | -Assembly of peat-based flowpot system is time consuming; Difficult to sample roots as they are stringly attached to peat substrates |
| Chesneau et al., 2025 <sup>24</sup> | MetaFlowTrain- Exometabolite mediated plant-microbe-microbe interactions | +Unidirectional flow of metabolites between organisms can be detected. |
|  |  | -Overlooks the complex feedbacks due to bidirectional exchange of metabolites between organisms |

### 1.4 Expertise needed to implement the protocol

Most steps involved in this protocol can be easily performed by a single graduate student or postdoctoral researcher with a basic understanding of plant tissue culture and molecular biology. However, microbiome composition and metabolomics analysis require high throughput infrastructure with trained personnel.

### 1.5 Limitations

Our system has certain limitations which arise primarily due to the nature of the set-up and to a certain extent, these limitations can be addressed through adoption of appropriate controls. We provide details on the current limitations of the system and the scope for addressing them:

a. *In-vitro system*: LEAP-CS is an *in-vitro* system to simulate phytobiome interactions, and, therefore, can be used to a limited extent to understand the phytobiome interactions under real cultivation regimes (soil/soilless). This system can serve as a quick screening tool to shortlist candidates or experimental designs for pot experiments or test bedding. Observations/results from this system could follow a similar trend as pot experiments or field trials, however, this would have to be established before drawing any extrapolations. *Suggestion to overcome limitation*: To make the system better, in place of conventional plant growth support media, alternative growth media enriched with soil extract, root exudate and microbial metabolites can be utilized.
b. *Microbial colonization*: LEAP-CS essentially mimics phytobiome interactions that happen in the rhizosphere region. Our system has a physical barrier between the host plants and the microbiome which does not allow for microbial colonization on the rhizoplane thereby, reducing the power to capture interactions in the endosphere or phyllosphere. *Suggestion to overcome limitation*: If experiment requires microbial colonization, then it can be done before seedlings are transplanted in the system (*via* root drenching) or it can be done before seeds are germinated (seed dressing). Alternatively, a system can be set-up without the membrane to facilitate microbiome colonization in host plants. However, this will require additional optimization and inoculum characterization to prevent system overload and contamination.
c. *Assay run time*: LEAP-CS utilizes membrane to facilitate physical separation between the host plants and the microbiome. Current membrane material starts to lose agility and strength if the system is run for longer than 10-15 days. However, certain host plants may require a longer run time depending upon their growth rate, environmental conditions, and experimental designs. *Suggestion to overcome limitation*: This limitation can be addressed by having them transplanted to the system closer to their growth-spurt or a critical stage depending on the experimental design. Alternatively, as suggested in the previous section, other materials can be explored which are of similar pore size and offer a physical separation between host plants and microbiome.
d. *Invasive sampling endpoints:* Sampling and harvest harvesting for multi-omics platforms and plant phenotyping is invasive in the current format for LEAP-CS. This would necessitate additional biological replicates and controls for a time-course based experiment where sampling is done at multiple time-points. This is a limitation as invasive sampling would increase the required resources including manpower to conduct a time-course experiment with multiple variables. *Suggestion to overcome limitation:* Monitoring of plant growth and phenotypic changes can be done non-invasively using the LEAP-live system in a semi-quantitative fashion. LEAP-live allows for real-time growth profiling by measurement of green pixels that corresponds to leaf growth. For other multi-omics analyses, plates must be sacrificed at each time-point for harvest in the current format.
e. *LEAP-CS is meant for early stage developmental spurts:* Current format supports seedling stage or small-sized plants due to the size limitations. Apart from the model species, other agronomically relevant plant species used for this system are short-cycle crops (35-45 days) where, seedling stage appears within 10-20 days of seed germination. *Suggestion to overcome limitation* Alternate formats can be adopted to suit the experimental needs for other host plants such as bigger plate sizes (as mentioned in the earlier section).

#### Inoculum optimization for different host plants

Interactions in the phytobiome, to a large extent, are influenced and determined by the host genotype, making it unique for each plant species. To study these interactions in LEAP-CS, the initial microbiome load in the system is critical, it should be high enough to facilitate active exchange of metabolites and growth of phytobiome, and optimum to avoid overwhelming the system. Hence, microbiome inoculum must be characterized before conducting an experiment and the starting inoculum requirements may differ based on experimental design and plant host in question.

## 2. Materials

### 2.1 Reagents

Reagents and biological materials for LEAP-CS set up
Seeds of Arabidopsis thaliana (wild type)-ARBC Stock
Compost soil (Jiffy Substrates, Smart)
PBS buffer (pH-7.4)
Murashige and Skoog basal salt mixture (Duchefa, cat. no. M0222)
Sucrose (Sigma, cat. no. 84100)
MES hydrate (Sigma, cat. no. M5287)
Sodium hydroxide (GCE Lab. Chem. cat. no. E7400)
Phytagel (Sigma, cat. no. P8169)
Bleach (Chlorox)
Tween 20 (Sigma, cat. no. P1379)

#### Reagents for microbiome analysis

Soil microbe DNA isolation Kit (Zymo,)
Ethanol Mol. Bio. Grade (Sigma,)

#### Reagents for analyses

Plant genomic isolation kit (Qiagen, cat. no. 69104)
Plant RNA isolation kit (Qiagen,)
Agarose (1^st^ BASE, cat. no. BIO-1000)
Tris base (1^st^ BASE, cat. no. BIO-1400)
Hydrochloric Acid (Brand, cat)
Acetic Acid (Brand, cat)
EDTA (Sigma, cat. no. ED2SS)
Floro Red Nucleic Acid Stain (1^st^ BASE, cat. no. BIO-5171)

### 2.2 Equipment

1.5/2 ml sterile safe-lock tube
15 and 50 ml sterile Falcon-tube
100 micron sterile Filter (Falcon, cat. no. 352360)
Table top centrifuge capable of holding 1.5-mL and 50 ml tubes (Thermo Fisher)
Vortexer
Laminar Air Flow
Vacuum Filtration Unit pore size 0.45 µm (Thermo)
Sterile square Petri plates (12 cm by 12 cm by 1.7 cm height)
Sterile Spreader
Parafilm
Pipette tips (20 μl; 200 μl; 1,000 μl)
Single-channel pipettes (2−20 μl; 20−2,00 μl; 100−1,000 μl)
Bench top pH meter
Cellulose dialysis membrane 14 kDa (Sigma, D9402)
Growth chamber or room with temperature, humidity and light control system
PCR machine (BioRad T100 Thermal Cycler)
Vertical Gel electrophoresis unit ()
Gel Documentation System (SYNGENE, G:BOX)
NanoDrop
Autoclave

### 2.3 Reagent set up

#### Primer design

Forward and reverse primers for microbiome composition study through 16S sequencing by IDT technologies

#### MS media

Mix Murashige& Skoog Basal Salt Mixture (company), 50 mM MES, 0.5 % weight/ volume of sucrose in MiliQ water. Adjust the pH to 5.8 using 5M Sodium hydroxide solution. Add 0.8% weight/ volume of Phytagel (Sigma-Aldrich) to prepare solid MS medium.

#### MS-agar Plate

Autoclave the MS medium with 0.8% phytagel. After that, aseptically pour 50 ml of the warm medium (60-65°C) to each square plate. Allow to solidify and store at 4 °C for future use.

#### Soil Extract Medium (SEM)

Take 70 gm of Jiffy compost soil in a 1000 ml screw-cap glass bottle and add MiliQ water to make the volume 1000 ml. Shake well and autoclave for 15 min at 15 lb pressure. After autoclaving again shake and keep at benchtop for overnight to cool down and to settle down the larger soil particles. After that, gently take out the extract from the top without disturbing the precipitates and pass through the vacuum filtration unit (pore size 0.45 µm). Add 1% weight/ volume of Phytagel (Sigma-Aldrich) to prepare solid SEM medium.

#### Seed sterilization solution

Prepare a solution of 50% chlorox and 0.2% Tween 20 for soaking the seeds for 5 min.

### 2.4 Equipment set up

#### Placing of membrane onto the lawn of microbiome in MS agar plate

The membrane is a dialysis tubing with molecular weight cut-off 14000 Da molecular weight cut-off: 14,000 Da, Average flat width: 76 mm (3.0 in.). The tubing is flat and clear, and provided in rolls. Cut the membrane according to the sections of MS agar plate. Soak the membrane in Mili-Q water for 5 min. Then gently separate the dialysis tube to get a two pieces membrane section from a single cut.

#### Microbiome Analysis

The collection of the microbiome from the LEAP plate has been detailed in the point number 45-46 of the Procedure section. DNA was extracted from the microbiome. Amplicon sequencing was performed for all the samples and downstream analysis was carried out by dada2 and phyloseq packages. The raw data has been submitted to the NCBI

#### Metabolomics analyses

All chromatographic separations were carried out on a Vanquish liquid chromatography (LC) system (Thermo Fisher Scientific, Waltham, MA, USA), using an ACQUITY Premier UPLC CSH C18 (2.1×100 mm, 1.7-micron) reversed-phase (RP) column (Waters, Milford, MA, USA). The column and auto-sampler temperature were set to 45 °C and 6 °C, respectively. The mobile phase used was ultrapure water with 0.1% formic acid (as solvent A) and LC-MS grade acetonitrile (ACN) with 0.1% formic acid (as solvent B) with the following linear gradient elution program: 0-1 min, 2% B; 1-11 min, 98% B; 11-12 min, 98% B; 12-12.1 min, 2% B; 12.1-16 min, 2% B. The flow rate was set to 0.3 mL/min, and the injection volume was 5 µL.

LTQ Orbitrap Velos Pro mass spectrometer (Thermo Fisher Scientific) was used as a high-resolution mass spectrometer (MS) for the profiling. Analyses were carried out in a mass range of 50-1000 m/z at a resolution of 60,000, with a source voltage of 4 kV (positive polarity mode). Capillary and source heater temperatures were maintained at 300 °C with sheath gas, auxiliary gas and sweep gas flow rate of 45, 15 and 1 (arbitrary units), respectively. Xcalibur v2.2 software package (Thermo Scientific Inc., Waltham, MA) was used for acquisition and preliminary raw data processing. The Raw data has been submitted to MetaboLights database.

Raw MS data were converted to. mzXML format using open-source Proteowizard software, then pre-processed by the open-source R-based XCMS software for molecular feature detection and alignment. Progenesis QI Ver. 2.6 (Waters) was further used for peak picking, chromatogram deconvolution, and identification of metabolites.

## 3. Procedure

The overall process of LEAP-CS is schematically demonstrated in Fig. 2. The process consists of pre-assembly and system assembly procedures with defined timelines as illustrated in Fig 2A.

**Figure 2:**
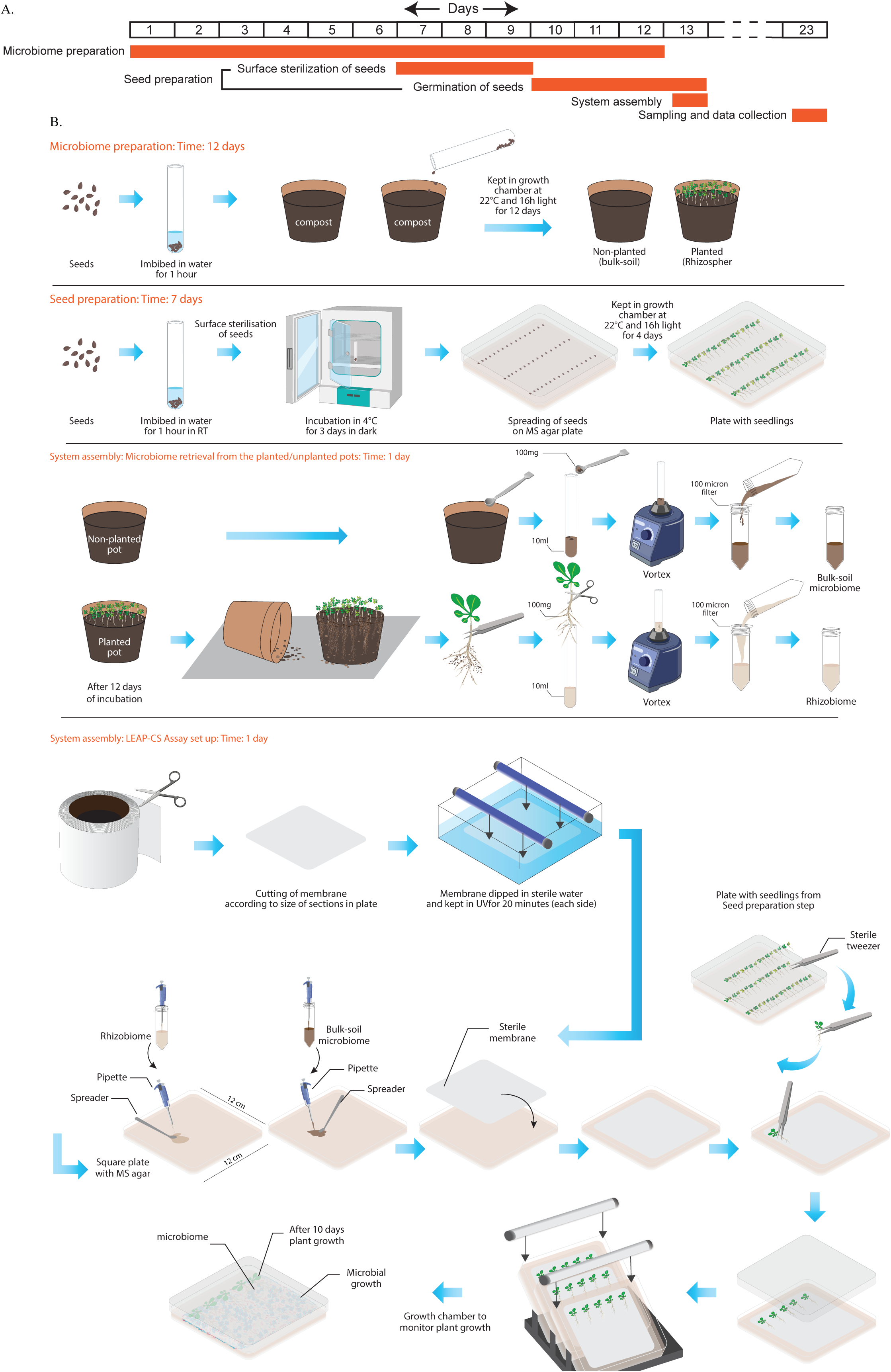
Diagrammatic Representation of the LEAP-CS assembly process. A. Time-line of the steps involved in LEAP-CS. B. Detailed assembly of components and workflow for setting up the LEAP-CS.

### 3.1 Pre-assembly procedures

The pre-assembly for LEAP-CS involves (i) preparation of microbiome inoculum; (ii) seed preparation involving seed treatment and germination.

#### 3.1.1 Preparation of microbiome inoculum: Timing 12 days

For purpose of demonstrating the system capabilities, native microbiome from the soil used to grow the model host plants (*Arabidopsis*) was used in LEAP-CS. This step entails preparation of native microbiome which can be adapted for other host plants as well depending upon the experimental design.

1. Take approximately 5mg of Arabidopsis seeds and imbibe in water for one hour.
2. Take two medium size pots and fill with approximately equal amounts of compost soil from the same source. CRITICAL STEP: Clean the pots properly and wipe with 70% alcohol before taking to avoid any microbial contaminations from pots.
3. Sow the imbibed seeds in one pot of compost soil and mark it as a planted pot. Mark another pot without seeds as a non-planted pot.
4. Keep both the pots in the growth chamber at 22°C with 16h light and 8 hrs dark cycle for 12 days.
5. Water both the pots at regular intervals in identical manner. Monitor the pots every day. CRITICAL STEP: Water the pots to only moist the soil. Excess watering may hamper the plant growth and microbial composition. ? TROUBLESHOOT

#### 3.1.2 Seed preparation: Timing 3 days

6. Take approximately 30 mg of *A. thaliana* seeds in a 2 ml centrifuge tube and imbibe for one hour at room temperature.
7. Centrifuge at 8000 x g for 15 sec. to settle down the seeds at the bottom of the centrifuge tube and gently discard the water.
8. Add 1ml of surface-sterilization solution (Material, section 2.3) to the tube and mix well with the imbibed seeds for 5 mins. CRITICAL STEP: Excess imbibition time in surface sterilisation solution may reduce the germination frequency of the seeds.
9. Centrifuge at 8000 x g for 15 sec. to settle down the seeds at the bottom of the centrifuge tube and gently discard the surface sterilisation solution.
10. Resuspend the seeds in 1ml of sterile MiliQ water within Laminar Air Flow (LAF).
11. Repeat the step ‘8’ for six times with the addition of 1ml of sterile MiliQ water each time. CRITICAL STEP: Any adherent of surface sterilisation solution can reduce the germination frequency.
12. Resuspend the seeds in 1ml of sterile MiliQ water and keep in dark at 4°C for three days for stratification.
13. After three days spread the resuspended seeds in three layers (each layer with 100 µl of resuspended seeds) on to the square plate with 0.8% agar (pH 5.8) containing MS media (4.4 g/L MS salt, 0.5 g/L 2-(N-morpholino) ethanesulfonic acid, and 5 g/L sucrose).
14. Keep the plates to germinate in a growth chamber at 22°C with 16 h of light for four days (Fig 2B). ? TROUBLESHOOT

### 3.2 System assembly: Timing 1 day

The assembly of the system was carried out in two phases performed on the same day. Details on the steps followed for system assembly are captured as a part of the illustration in Fig 2B.

#### 3.2.1 Phase-I-Microbiome retrieval from the planted/unplanted pots

Microbiome was retrieved from the twelve days pots as discussed in earlier section 3.1.1. Microbiomes from the rhizosphere of the planted pots were designated as rhizospheric and unplanted pots were designated as bulk-soil microbiome respectively. Standard documented procedures were followed to harvest the microbiome, details about the procedures are provided below.

15. Take 10 ml of liquid SEM medium in a sterile 15 ml falcon tube and weight it in a weighing machine.
16. Place the soil from the planted and non-planted pots separately onto a sterile aluminium foil under aseptic conditions in the LAF.
17. Remove the plants from the soil using sterile tweezers carefully to ensure root integrity.
18. Remove the excess soil particles from the roots carefully (Fig 2B: System assembly, Supp Fig 1a). CRITICAL POINT: keep the layer of soil particles attached with the roots.
19. Detach the roots with attached soil carefully and transfer into the 10 ml of liquid SEM medium.
20. Weigh the falcon after putting the roots with soil particles upto 100 mg and mark that as rhizospheric microbiome.
21. Similarly take 100 mg soil from the non-planted pots in 10 ml of liquid medium in another falcon tube and mark that as bulk-soil microbiome. CRITICAL STEP: Precision in weighing is required to quantify the microbiome density.
22. Vortex both the tube for 3 times each with 1 min to detach microorganisms from the soil.
23. Place a sterile 100 micron filter on a 50 ml falcon tube separately for rhizospheric and bulk-soil microbiome within LAF (Supp fig 1b).
24. Filter the respective microbiome suspension to the specific tube.
25. Use the filtrate from the respective tube as seed microbiome or inoculum for rhizoshpere and bulk-soil.

#### 3.2.2 Phase-II-LEAP-CS Assay set up

This is the final step in the assembly of the LEAP-CS to assess the plant-microbe interaction and its effects on the growth of plants. The procedure to set up the assay is described as follows:

26. Take square sterile petriplates with 50 ml of SEM-Agar.
27. Cut the 0.2μm dialysis tubing cellulose membrane (Sigma-Aldrich) with a molecular weight cut off 14,000 Da according to the size of in the plates.
28. Soak the pieces of membrane in a MiliQ water for 5 minutes. CRITICAL STEP: Soaking is necessary to separate the membrane.
29. Pour small amount of sterile MiliQ water into a sterile blank periplates within LAF in a sterile condition.
30. Place each piece of membrane onto the water layer in the petriplates within the LAF. CRITICAL STEP: Do not let membrane dry because drying will change membrane texture and property.
31. Switch on the UV light of the LAF for 20 min to disinfect the upper layer of the membrane.
32. After 20 minutes, flip the membrane to expose the bottom surface under UV light again for 20 min.
33. Take 100μl of each microbiome (rhizospheric and bulk-soil) and spread onto the medium in the plates in a sterile condition within LAF. CRITICAL STEP: Strictly avoid cross-contamination between rhizobiome and bulk-soil microbiome within the plate during placing of the membrane onto each section of microbiome layer.
34. Take the plates with 4 days old Arabidopsis seedlings (Step-12) from the growth chamber. ? TROUBLESHOOT
35. Gently place the seedlings with a sterile tweezer onto the membrane of each section of the plate within LAF.
36. Cover the plates with lids and seal with parafilm.
37. Keep the plates in growth chamber at 22°C with 16 h of light in vertically inclined position for monitoring, sampling and data collection upto 15 days. ? TROUBLESHOOTS

### 3.3 Sampling and data collection: Timing 10-15 day

38. Monitor the growth of plants and microbes in the plates daily and document as images.
39. Try to find out any noticeable growth difference observed between the rhizospheric, bulk-soil and No microbiome associated plants within the plates. CRITICAL STEPS: The time points of the observable difference may vary with plant types. In case of Arabidopsis the difference observed within 8-10 days in LEAP-CS.
40. Chose time points of sampling from the LEAP-CS plates based on the observed phenotypes and research hypothesis. CRITICAL STEP: Process of sampling in this system is destructive. So prepare as many as replicate to study any dynamics.
41. At the selected time point, open the plate within LAF and gently detach the plants from the membrane.
42. Wash the plants roots in sterile MiliQ water for metabolomics study of root exudates.
43. Process the plants for further molecular study. PAUSE POINTS: Store the plants suitably in -80°C for future use
44. Gently remove the membrane from the media plates separately using sterile forceps and wash in sterile MiliQ water for metabolomics study to detect membrane-associated exudates. CRITICAL STEPS: The membrane is hard to remove from the agar layer as it become fragile with time in the LEAP-CS. It is recommended to finish the experiment and do sampling within 15 days of the assay set up.
45. 5 and 10 mL of sterile ultrapure water (RT), were added to the root and membrane respectively, and incubated for one hour under continuous mild shaking (130 rpm). The collected exudates were then centrifuged at 8,000 g at 4 °C for 10 min, filtered by 0.22-um PES syringe filters, and stored at -80 °C. They were subsequently freeze-dried and reconstituted in 200 µL of 50% methanol, and kept at -80 °C prior to analysis.
46. Scrape the microbiome of each section separately into sterile sterile ultrapure water for further composition and molecular study. PAUSE POINTS: Store the microbiome suitably in -80°C for future use.
47. The microbiome layer was resuspended in 3 mL of sterile ultrapure water (RT) from the plates and incubated for one hour under continuous mild shaking (130 rpm). The sample was then centrifuged at 8,000 g at 4 °C for 10 min. The upper supernatant was collected and filtered by 0.22-um PES syringe filters, and stored at -80 °C. They were subsequently freeze-dried and reconstituted in 200 µL of 50% methanol, and kept at -80 °C prior to analysis. The microbial pellet was snap was snap-frozen in liquid nitrogen and kept at -80 °C for composition analysis.

#### Troubleshooting

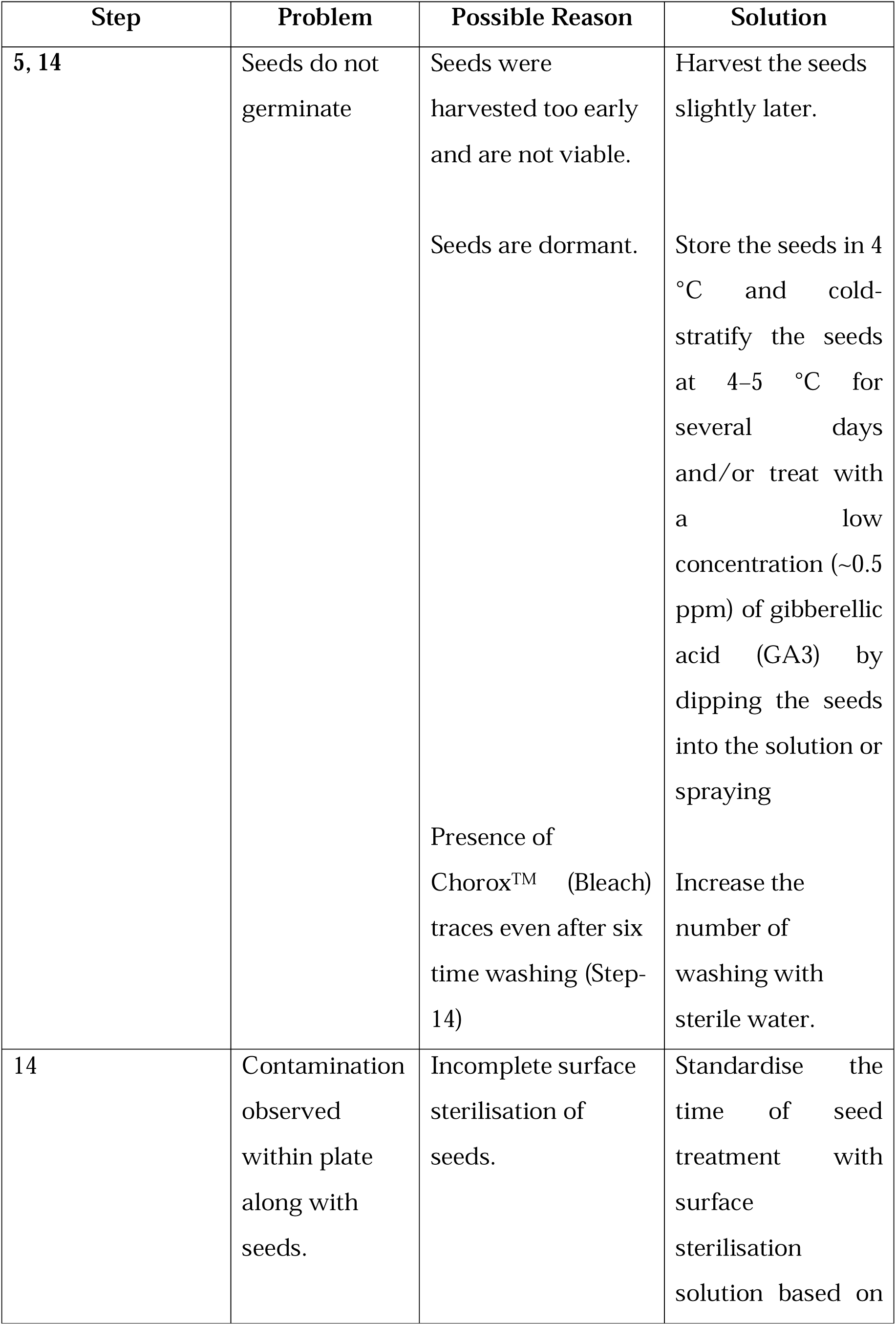
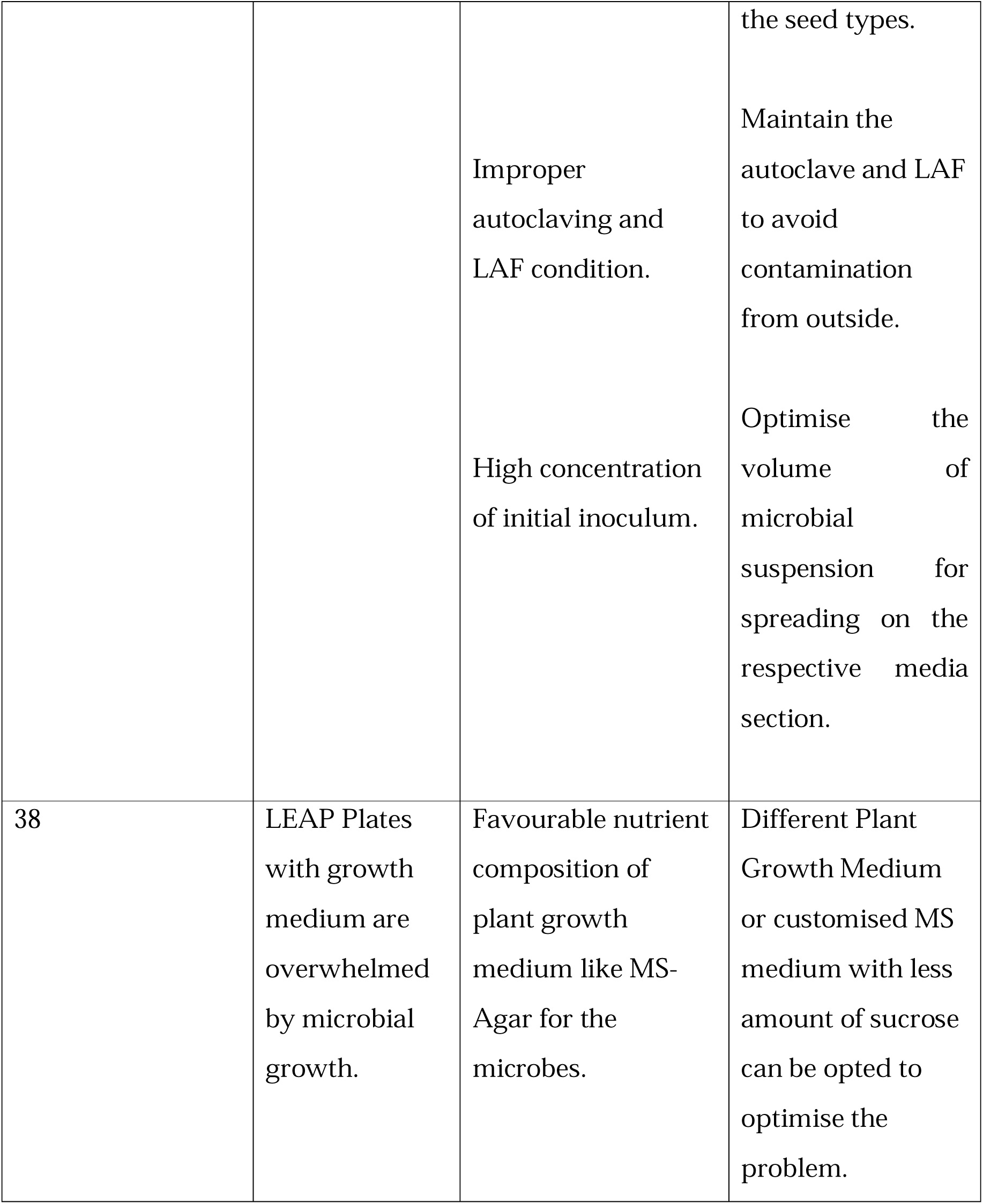

## 4. Anticipated results

The LEAP-CS system can be used to investigate both plant and microbiome interactions in a controlled environment. Microbiome inocula prepared from different sources, such as rhizosphere or bulk soil, can be used in the system. Soil extract medium facilitated recapitulation of some of the chemical diversity in the natural system for investigating plant-microbe interactions. The use of membrane allowed small molecules to be exchanged between the plants and microbiome. We used LEAP-CS to obtain data on plant growth dynamics, metabolic features of root exudates, and microbiome composition. Axenic 4 days old Arabidopsis seedlings were grown in the system with bulk-soil, rhizospheric, or no microbiome over 15 days in a growth room with a controlled environment (Fig. 2B). Several studies were conducted to determine the functional impact of microbiota on the plants and microbiome composition, as well as the exometabolome released by either roots or microbes. (Fig. 3e).

a. Plant growth dynamics: Plant growth was monitored in a non-destructive manner using both low-cost RGB cameras and the WinRHIZO scanner at regular intervals up to the 13th day. Representative images of plants grown with bulk-soil, rhizospheric, and no microbiome associations are shown in Fig. 3a. Quantitative measurements of plant growth, specifically leaf area, clearly indicated greater growth in the bulk-soil condition compared to the rhizospheric and no microbiome conditions (Fig. 3b).
b. Microbiome analysis: After 13 days of plant-microbiome association in the LEAP system, the set-up was dismantled, and the plants were set aside to isolate root exudates and other molecular studies. The composition of the microbiome below the membrane was determined using the 16S rRNA amplicon sequencing. The LEAP system supported the growth of approximately 146 ASVs distributed across 10 phyla, making this microbiome moderately complex, compared to that of the soil. As expected, the inoculum samples for both bulk-soil and rhizospheric microbiomes showed the highest alpha diversity compared to the respective LEAP plate isolates (Fig. 3d, Supp table 1).
c. Metabolome analysis: We performed untargeted exometabolome analysis on the root (Root Exometabolites, RM), membrane (MM), and microbiome (Microbiome Exometabolites, MIM) sections of LEAP plates inoculated with either bulk or rhizospheric microbiomes. Chromatographic techniques coupled with high-resolution liquid chromatography–mass spectrometry (LC-MS) were employed to obtain high-quality mass spectra for metabolite identification and classification.

**Figure 3:**
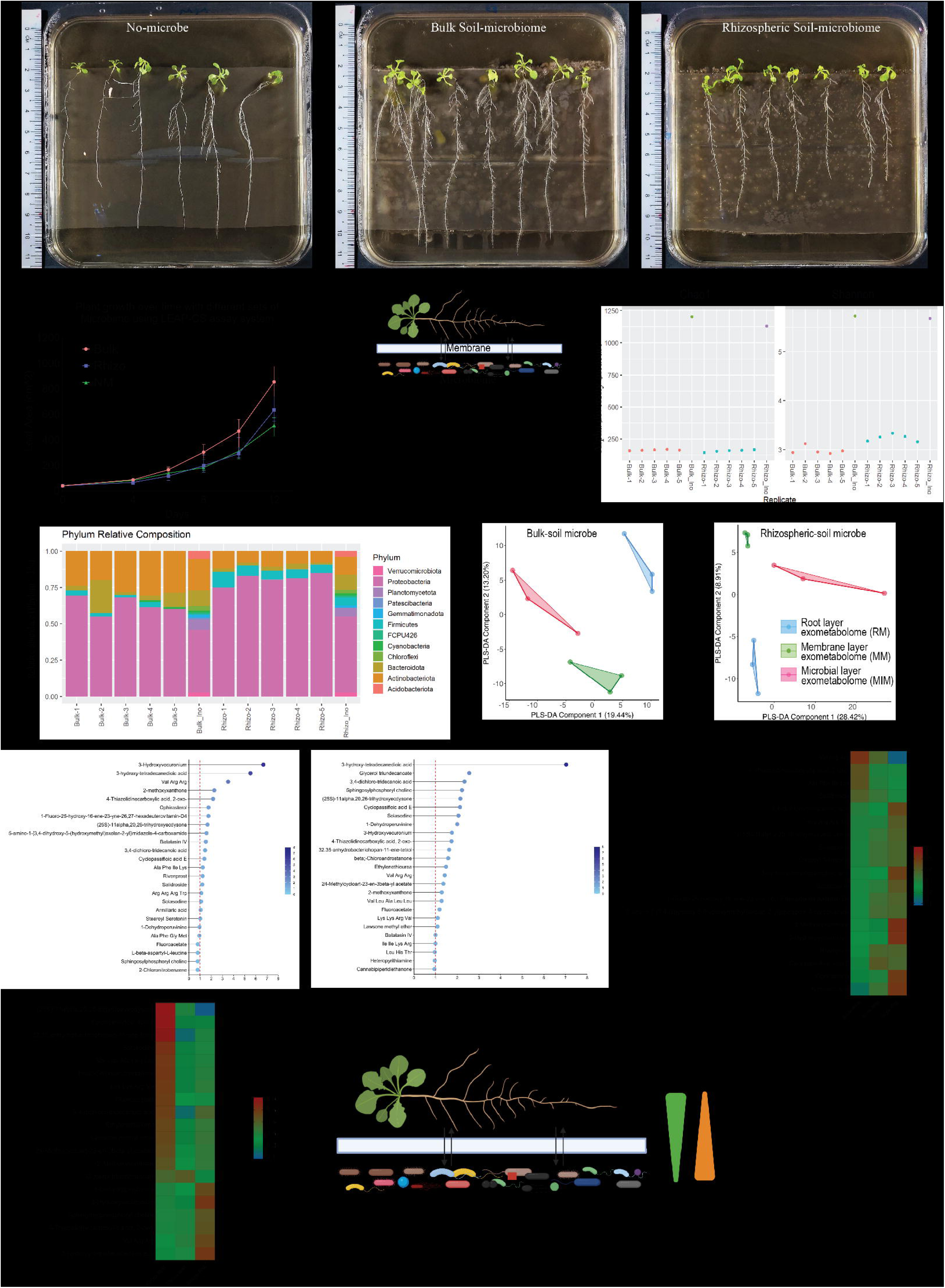
Fig 3: Anticipated results from LEAP-CS: a: Phenotypic changes of Arabidopsis grown with microbiomes from the bulk or rhizosphere region, compared to the control with no microbiome inoculum b: Line graph showing the growth dynamics of plants associated with bulk (orange), rhizospheric (blue), and no microbiome (green), as indicated by non-destructive leaf area measurements over 12 days (n = 5). c: Workflow for microbiome and metabolome studies from LEAP-CS plates from deferent section. d. Alpha diversity of microbiome from LEAP plates. e. Microbial compositions from bulk and rhizosphere compartments after 12 days of growth on LEAP-CS plates. f: The PLS-DA plot of the metabolic features of the exometabolome from Root (RM), Membrane (MM) and Microbial (MIM) section of LEAP-CS plates with bulk and rizospheric microbiome (n=3). g, h: The VIP score plot of the annotated features with their abundance in different section of LEAP plates with bulk (g) and rhizospheric (h) soil microbiome. i, j: Heat-map showing the abundances of the annotated compounds having VIP score>1 across the different sections, from RM to MIM, within a given LEAP plate with bulk (i) and rhizospheric (j) soil microbiome. k. The capability of LEAP-CS to assume the flow of metabolites across different sections like RM, MM and MIM.

In total around 688 metabolic features have been identified from the LEAP plates inoculated with either bulk or rhizospheric soil microbiomes (Supp table 2). These data are available for mapping to metabolic pathways using public platforms such as MetaboAnalyst, MAPPS, and others. Partial least squares discriminant analysis (PLS-DA) of the metabolic features from the RM, MM, and MIM showed clear group separation in both the bulk and rhizospheric microbiome LEAP setups (Fig. 3f). Variations of the metabolic profiles along components 1 and 2 as showed in the PLS-DA plots indicate metabolite dissimilarities across different sections, from RM to MIM, within a given LEAP plate. Analysis of the variable importance in projection (VIP) scores following metabolite annotation (Supp table 3)identified key metabolites contributing to these profile variations in both bulk and rhizo LEAP set up (Fig. 3g, h). Metabolites with VIP scores greater than 1 showed a distinct gradient across sections, suggesting their probable origin and potential transport pathways as the source sink relation. For example, compounds that are more abundant in a specific section are likely to originate from that section. Several metabolites were found to be more abundant in either the root or microbial exometabolomes, indicating their respective sources and sink (Fig. 3i, j).

This system enables the study of changes in key metabolites, such as signaling molecules, alongside associated shifts in microbiome composition (Fig. 3k). Thus, it provides a valuable platform for elucidating the chemical basis of plant–microbiome interactions.

### Potential Application of the LEAP system

LEAP-CS is designed to characterize physiological, molecular, and metabolic phenotypes to understand (i) plant-microbe and microbe-microbe interactions; (ii) metabolic exchanges between the plant and microbes with the prediction of their probable source and crosstalk; (iii) microbial community shifts across different time and growth phases. This system allows simultaneous retrieval or collection of plants, root exudates and microbiomes at the end of the assay. Plant phenotyping, root exudates profiling and microbiome analysis can be performed from the same LEAP-CS plates (Fig 3). The quality and amounts of DNA and RNA recovered from plants and microbiome module are adequate to perform 16S rRNA sequencing, metagenomics and metatranscriptomics analyses to understand the community composition, its potential and realized function (Fig 3). This system can also be used for quick lab-based screening of several microbes or microbial consortia to determine their impact on plant growth. This LEAP system provides the researchers with a tool to address several fundamental and translational research aspects, especially in the field of plant-microbe interactions.

### Applications for basic science studies

LEAP-TOL: Effects of biotic and abiotic stresses on plant-phytobiome interaction can be evaluated using suitable supplements in the medium. (Fig 4). Solid support media composition can be manipulated to mimic abiotic stress such as through addition of osmotically-active mannitol or polyethylene glycol (PEG) to create drought stress. Similarly, a pathogen can be introduced in the system at the right stage along with beneficial microbiomes to trigger biotic stress in the system.

**Figure 4:**
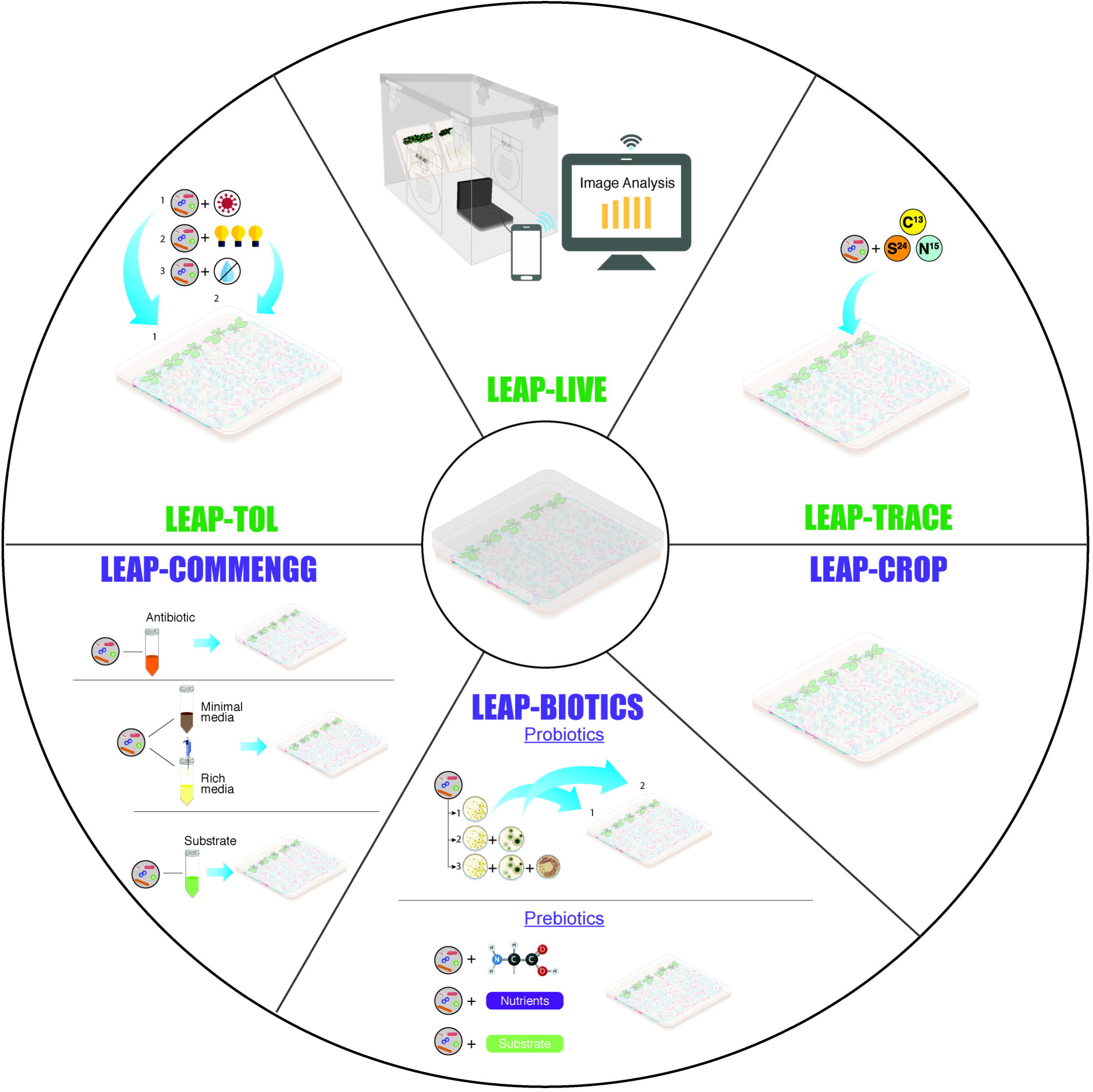
Potential applications of LEAP-CS for basic science studies and translational applications. LEAP-CS is versatile and modular in nature. It can be set up with minimal modifications for different research objectives. Potential LEAP-CS extensions to study (i) growth profiling in real time; (ii) plant volatilome as influenced by specific microbes/microbiome; (iii) effects of abiotic and biotic stress on plant-microbiome/microbe interactions to answer fundamental research questions. For translational applications, the LEAP-CS can be modified to (i) perform microbiome manipulations and engineering and (ii) screen potential microbial isolates for synthetic consortia development (iii) expand its scope to include economically important crops for agricultural applications.

LEAP-TRACE: To further investigate the chemical interactions and metabolic exchanges between the phytobiomes, controlled tracer studies can be conducted in LEAP-CS and labelled substrates can be introduced either through the support media for host plants or by enriching microbiomes in media containing stable isotopically-labelled substrates depending upon the experimental design (Fig 4). This module can be useful to study nutrient acquisitions and transformations in the phytobiome, which could offer insights for nutritional provisioning by microbiomes.

LEAP-LIVE: One of the potential fundamental applications includes real-time growth profiling and plant phenotyping. This system can be used to design a non-invasive, real-time growth profiling of plants and can be integrated with image processing software to perform plant phenotyping.

### Translational

LEAP-CROP: Small sized crop plants such as leafy vegetables, medicinal or ornamental plants can be grown in the presence of phytobiomes to uncover host-microbe interactions relevant to agronomic traits.

LEAP-BIOTICS: LEAP-CS can be utilized to perform screening of plant growth promoting microbial strains for development of agricultural probiotics and synthetic microbial consortia (Fig 4). As the system is relatively simple, it offers a unique platform to screen microbial isolates in different combinations to study their effects on plant hosts and their interactions. This can be particularly helpful in shortlisting candidates/combinations for pot experiments that usually precedes test-bedding and field trials. This application can also be extended to screen prebiotics or substrates that enhance stability and efficiency of microbial probiotics.

LEAP-COMMENG: Microbiome module in the system can be selectively enriched before plating it on the assay plate. The selective enrichment of microbial communities facilitates microbiome engineering for better plant host phenotypes. It further assists in studying the internal dynamics between the microbial communities which is instrumental for development of stable microbial consortia.

## Acknowledgements

The authors thank the NUS, NERI for providing the facility of metabolite analysis. We are grateful to Dr. Gourvendu Saxena and Dr. Terence Wong for transferring their previous knowledge to successfully designing of the protocol.

## Conflict of Interest Statement

The authors do not have any competing interest.

## Author Contributions

S.S conceived and supervised the project, Manuscript writing; M.M and S.P Experimental Design and Manuscript writing; M.M Wet lab experiments, plant phenotype data collections, 16S rRNA amplicon and metabolite sample preparation, figure generation; R.B, A.M and S.M.M metabolite data analysis, figure generation and manuscript editing; I.C.H.T prepared the infographics related to this manuscript.

## Funding

This work was supported primarily by the Singapore Centre for Environmental Life Sciences Engineering (SCELSE).

---

**Supplementary Figure 1:**

Preparation of microbial inoculum for LEAP-CS mesocosm.

**Supplementary Table 1:**

ASV table from 16S rRNA amplicon sequencing data with their abundance. The whole file contains several sheets for ASV number with their abundance, taxonomic identification and sample meta data information.

**Supplementary Table 2:**

Feature table from the metabolic analysis from the LEAP-CS plate. The whole file contains several sheets for feature number with their abundance in different section of the LEAP plate, m/z value correspond to each feature and sample meta data information.

**Supplementary Table 3:**

Annotation of the compounds from the features

## Notes

### Competing Interest Statement

The authors have declared no competing interest.

### Summary of Updates

Figure numbering updated. In the earlier submission the figure 4 appeared as figure 3 and figure 3 appeared as figure 4. In this submission, the numbering has been updated.

