## Supplementary material for "Live-Exudation Assisted Phytobiome Cultromics System (LEAP-CS): A High-Throughput Cultromics System for Studying Plant-Microbiome Interactions through Diffusible Metabolic Exchange": Supp Fig 1

a. b.

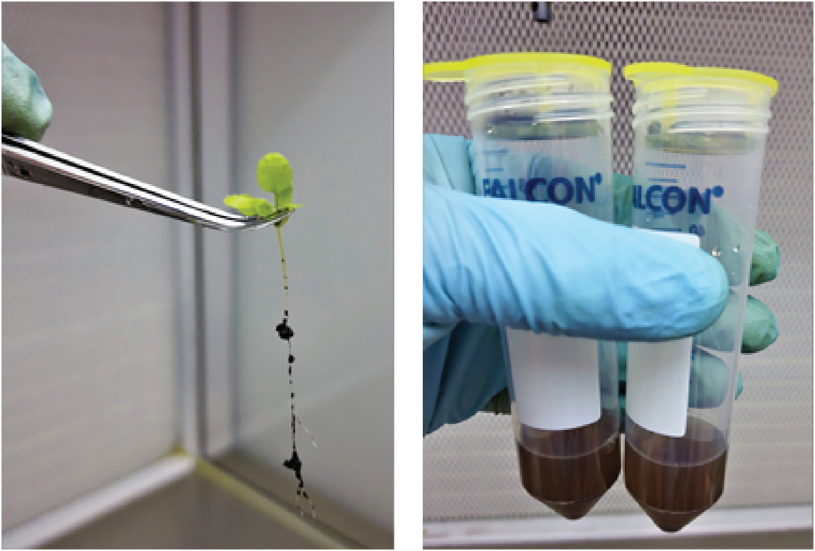

4 days old seedling with rhizospheric soil attached with roots

Microbiome suspension from bulk and rhizospheric soil
